# An adaptive control scheme for Interleukin-2 therapy

**DOI:** 10.1101/2020.02.21.959221

**Authors:** Sahamoddin Khailaie, Ghazal Montaseri, Michael Meyer-Hermann

## Abstract

Regulatory T cells (Treg) are suppressor cells that control self-reactive and excessive effector conventional helper T cell (Tconv) responses. Breakdown of the balance between Tregs and Tconvs is a hallmark of autoimmune and inflammatory diseases. Due to the positive dependency of both populations on Interleukin-2 (IL-2), it is subtle leverage to restore the healthy immune balance. By employing a mechanistic mathematical model, we studied the IL-2 therapy in order to increase and stabilize Treg population and restrict inflammatory Tconv response. We introduced an adaptive control strategy to design the minimal IL-2 dosage. This adaptive strategy allows for an individualized therapy based on the feedback of immune kinetics of the patient. Our *in silico* results suggest that a minimal Treg population is required to restrict the transient side-effect of IL-2 injections on the effector Tconv response. The combination of IL-2 and adoptive Treg transfer therapies is able to limit this side effect in our simulations. Implications of our *in silico* results are discussed in the context of autoimmunity and transplantation.

## Introduction

Among numerous factors that control immunological tolerance, the balance between regulatory T cells (Treg) and conventional T helper cells (Tconv) is indispensable. By suppressing Tconv through various mechanisms (1), Tregs can control magnitude and duration of inflammatory responses in order to maintain healthy immune homeostasis and protect the host from immune-mediated pathology. Manipulation of homeostasis and interplay of Treg and Tconv is a therapeutic approach in the context of autoimmunity, transplantation, and cancer where inflammation works against the patient. In cancer, enhanced Treg homeostasis is deleterious, while quantitative and qualitative defects in the Treg compartment are implicated in multiple autoimmune diseases in humans and mice (2).

In the absence of foreign antigens (Ag), the homeostatic number of Tregs and Tconvs is under control of homeostatic proliferation and thymic export. Activation of both, Treg and Tconv, via their T cell receptor (TCR) is stimulated by antigen presenting cells (APCs) (3, 4). Interleukin-2 (IL-2), a monomeric glycoprotein, is characterized as a proinflammatory cytokine that is predominantly produced by activated Tconvs, but not Tregs. IL-2 acts as an autocrine T cell growth factor and is necessary for the survival of Tregs and proliferation of both Tregs and Tconvs (5–8). Given that the biological factors impacting survival, activation, and proliferation of Tregs and Tconvs are mostly shared, immunotherapeutic perturbations using such factors impact both populations and demand caution. For example, with the premise that IL-2 stimulates the effector T cell population, high dose of IL-2 administration were extensively used in cancer patients, despite poor safety profile due to side effects (9). The therapeutic outcome was only partially successful and the unintended impact of IL-2 on Treg expansion, at least in part, may explain the failure of this therapy in some cancer patients (10–13).

Benefits of low-dose IL-2 administration were shown in patients with HCV-induced vasculitis (14), type 1 diabetes (15), and chronic graft-versus-host disease (GVDH) (16), which resulted in significant Treg proliferation and resolving the deficiency of Treg numbers in the context of autoimmune and alloimmune inflammatory diseases. Such results motivated search for optimal doses and frequencies of IL-2 administration (15, 17, 18).

Due to the subtle role of IL-2 in regulating both effector and suppressor arms of the immune system, IL-2 therapy could act like a double-edged sword and result in unintended adverse outcomes. Mathematical modeling is a candidate approach to explore IL-2 therapy. In this study, by employing a mechanistic mathematical model of T cell responses, we conducted an *in silico* analysis of IL-2 therapy as an approach to increase and stabilize the size of either T cell subset. We introduced a feedback control scheme to calculate a time-resolved adaptive IL-2 dosing for each individual *in silico* patient. This scheme is based on the “impulsive zone model predictive control (iZMPC)” algorithm (19, 20). The adaptive algorithm calculates proper IL-2 doses at each injection episode based on (1) feedback from the current status of the patient’s immune response, (2) prediction of how the immune response progresses according to the mathematical model for a limited time horizon, and (3) predefined desired range and constraints for the immune variables (Tconvs, Tregs and IL-2). We provided qualitative results as a proof-of-principle for our methodology in the context of transplantation as a use-case scenario. In this scenario, restricting acute/chronic effector responses against the graft by increasing and stabilizing Treg numbers is the main goal. But the adaptive IL-2 dosing algorithm is general and can be employed in other inflammatory contexts, such as cancer where increasing the number of effector T cells is desired.

## Materials and Methods

### Mathematical description of T cell responses

We described the dynamic interplay of activated Tconv (T) and Treg (R) populations under the influence of IL-2 (I) as depicted in **Figure 1** by the following set of ordinary differential equations (ODE) (21)

**Figure 1:**
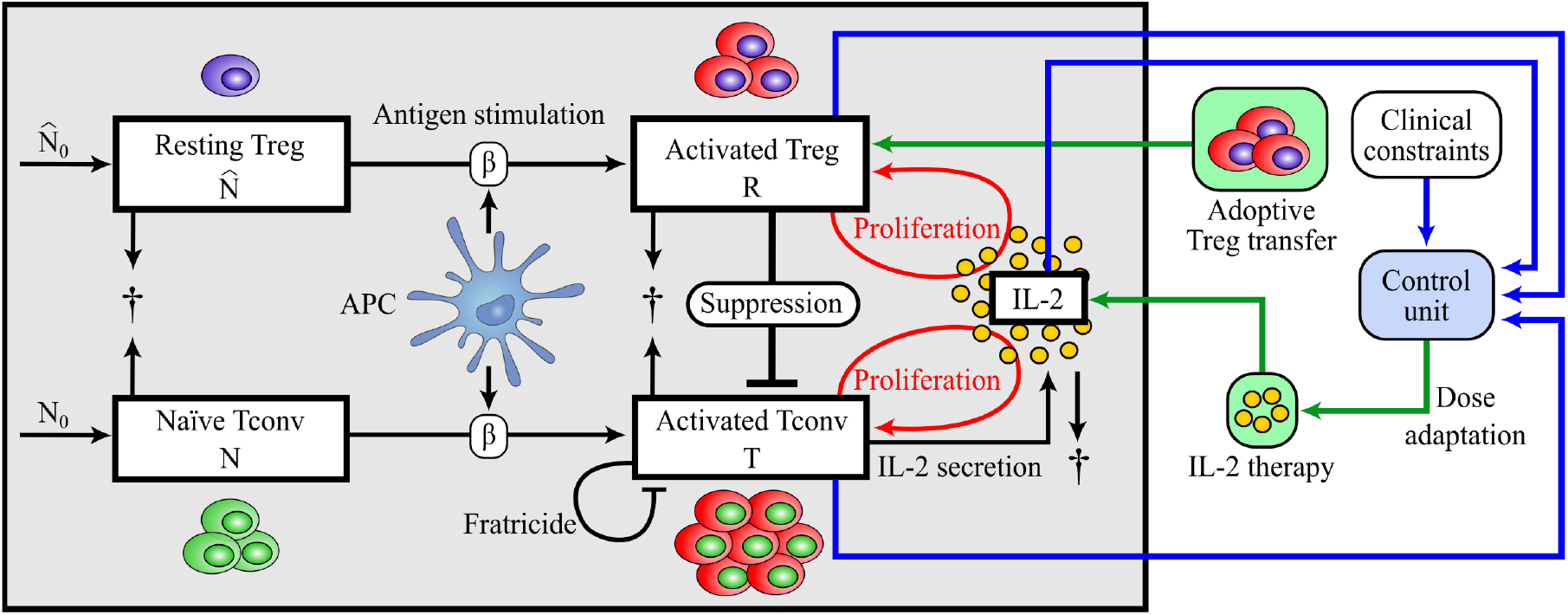
Scheme of the T cell response model and adaptive dosing IL-2 therapy. IL-2 concentration and the population of two T cell subsets, Tconvs and Tregs, are the immune variables considered in the mathematical model (1). Naïve Tconvs and resting Tregs originated from thymic selection are under homeostatic turn-over in the periphery. Upon Ag-stimulation provided by APCs, naïve Tconvs and resting Tregs become activated. In contrast to activated T cells, activated Tregs do not secrete IL-2, but both activated populations proliferate in dependence on the presence of IL-2. Activated Tregs suppress activated Tconvs in a cell-contact dependent and cytokine-driven manner. In contrast to Tregs, activated T cells undergo Fas-induced apoptosis by interacting with each other (fratricide). All cells undergo natural cell death and IL-2 is degraded. In the context of IL-2 therapy, the control unit provides the next optimal IL-2 dose according to a feedback from the current status of the immune variables. The control unit calculates the IL-2 dose that is needed to keep T cell numbers and systemic IL-2 concentration in a pre-defined range (clinical constraints). Adoptive Treg transfer is the therapeutic process of increasing Treg numbers in the immune system by transiently transferring Tregs to the individual.

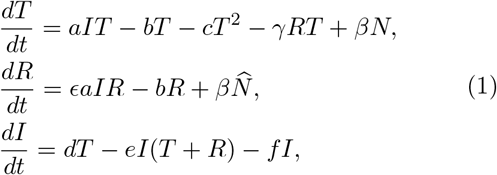

where *N* and 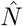 denote naïve T cells and resting Tregs, respectively, and follow

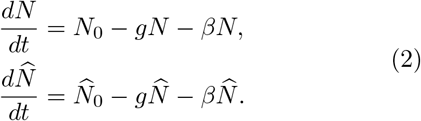

The definition and values of parameters are given in **Table 1**. We assume that activation of Tconvs and Tregs in the presence of sufficiently strong Ag-stimulation (large *β*) occurs much faster than other processes such as proliferation. Then, the variables *N* and 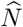 have faster dynamics compared to *T*, *R* and *I*, and can be treated in the quasi-steady-state approximation 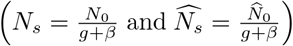, by replacing *N* and 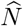 with *N*_*s*_ and 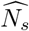, respectively. The parameter values of the model are tuned to mathematically obtained regimens where T cell dynamics show qualitative similarities to the experimentally observed T cell kinetics. Therefore, numerically obtained variables and times are given in arbitrary units, and the results shall be interpreted qualitatively. The parameter tuning process is based on a hierarchical modeling approach described in (21).

**Table 1:** Parameters of immune activation model.

| Parameter | Value | Description | Dimension |
| --- | --- | --- | --- |
| $a$ | 0.4 | Proliferation rate of activated T cells | molecule<sup>-1</sup>time<sup>-1</sup> |
| $b$ | 0.1 | Natural death rate of activated T cells and Tregs | time<sup>-1</sup> |
| $c$ | 10<sup>-5</sup> | Fratricide death rate of activated T cells | cell<sup>-1</sup>time<sup>-1</sup> |
| $d$ | 0.01 | IL-2 secretion rate by activated T cells | molecule<br>cell<sup>-1</sup>time<sup>-1</sup> |
| $e$ | 0.01 | IL-2 consumption rate by activated T cells and Tregs | cell<sup>-1</sup>time<sup>-1</sup> |
| $f$ | 1 | IL-2 decay rate | time<sup>-1</sup> |
| $g$ | 0.1 | Natural death rate of naïve T cells and resting Tregs | time<sup>-1</sup> |
| $\beta$ | 0.05 | Ag-stimulation of naïve T cells and resting Tregs | time<sup>-1</sup> |
| $\gamma$ | 0.1 | Treg-mediated suppression rate | cell<sup>-1</sup>time<sup>-1</sup> |
| $\epsilon$ | 0.6 | Ratio of proliferation rates of Tregs to Tconvs | - |
| $\lambda$ | 0.006 | Ratio of renewal rates of resting Tregs to naïve T cells | - |
| $N_0$ | 4 | Renewal rate of naïve T cells | cell time<sup>-1</sup> |
| $\hat{N}_0$ | $\lambda N_0$ | Renewal rate of resting Tregs | cell time<sup>-1</sup> |

### Mathematical description of IL-2 therapy

After intravenous administration of a drug, instantaneous jumps are observed in the drug concentration in plasma and target organ (22). In the framework of mathematical modeling, systems with discontinuities in their dynamics can be categorized as impulsive systems and allows for application of control engineering design tools (19, 20, 23–31). Among these methods, model predictive control (MPC) techniques have been widely used (19, 20, 23–28) due to their ability to consider systemic constraints, which confine the dynamics of the system variables or the external control input. In the context of pharmacodynamics, the control input is the administered drug.

MPC algorithms follow a receding horizon strategy to construct a feedback control law, here, the amount of the administered drug. At each administration time, system variables are measured and provided to the MPC algorithm. Given the horizon parameter *N*, MPC predicts the system dynamics for the next *N* steps using the current measurement as the initial value of the dynamics. Then, it calculates the next *N* optimal doses by solving a constrained optimization problem aiming at steering the system dynamics to the desired and predefined equilibrium points. After calculation of the *N* optimal doses, only the first dose is kept and administered. This process is repeated at each next drug administration time point. In this study, we employed the iZMPC algorithm (19, 20), an advanced version of MPC, in which system dynamics are moved toward a desired equilibrium set or space (instead to an equilibrium point) no matter which point inside the set.

Suppose an IL-2 administration scheme where IL-2 doses are sequentially injected intravenously at time intervals *τ*_*i*_, where *i* is an increasing sequence of positive integers. We assume equidistant IL-2 injection times, i.e., *τ*_*i*+1_ – *τ*_*i*_ = *δ*. For *t* ≠ *τ*_*i*_, variables *T* (*t*), *R*(*t*) and *I*(*t*) follow equation (1). At the moment of IL-2 injection (*τ*_*i*_), we assume a sudden change in the amount of IL-2, i.e.

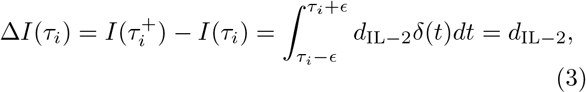

where 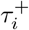 denotes the time instance after *τ*_*i*_, *δ*(*t*) is Dirac delta function, *ϵ* → 0, and *d*_IL−2_ is the dose of IL-2 injection. Considering the mathematical description of T cell responses under IL-2 therapy **Figure 1**, the problem of finding adaptive doses is formulated as the calculation of *d*_IL−2_.

By employing iZMPC, the optimal IL-2 doses are obtained by repeatedly solving a constrained optimization problem at each injection time using the current measurement of the system dynamics *T*, *R* and *I*. During the treatment, the amount of *T*, *R* and *I* should be kept in the allowed physiological ranges 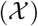. To have a successful treatment, system dynamics should remain in the therapeutic target window 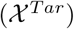. In addition, the injected dose should be confined to a range 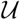 predefined by the safety or toxicology limitations. The physiological and therapeutic ranges as well as the safety considerations are implemented in the iZMPC and the optimal dose is calculated if the optimization problem is feasible. Infeasible cases can be solved by enlarging the constraining ranges or reducing the drug administration time interval. For more details about the iZMPC algorithm and the inclusion of design constraints, see the Supplemental Information (SI).

## Results

### Dynamics of T cell responses under chronic Ag-stimulation

To evaluate the effect of IL-2 therapy on the dynamics of activated Tregs and Tconvs (the term “activated” is omitted hereinafter), a scenario without therapy is considered at which the T cell response model (1) is stimulated with a sufficiently high chronic Ag-stimulation (a high constant value for *β*) such that a substantial proliferative Tconv response is initiated. It is assumed that the model starts from a healthy initial condition, i.e. all cells are in non-activated state. This scenario could represent the immune challenge in transplantation where the Ag-stimulation starts and stays chronic after the surgery.

In the absence of IL-2 therapy, the interplay of Tregs and Tconvs results in an oscillatory response (limit cycles) in all variables (see **Figure 2**). The initial (acute) response of Tconvs consists of an initial rise that resulted from the activation of naïve Tconvs by Ag-stimulation as well as the proliferation associated with the secreted IL-2. Tregs that are activated with the same Ag-stimulation cannot efficiently proliferate and suppress Tconvs until the concentration of IL-2 secreted by Tconvs rises. When Tregs proliferate, the number of Tconvs declines due to the direct suppression as well as IL-2 consumption by Tregs. The delay between the peaks of Tconvs and Tregs in the model results from the dependence of Tregs on IL-2 for proliferation and their inability to secrete this growth factor. The persistent stimulation of T cells with Ag (constant *β*) is responsible for re-initiation of another Tconv responses after having been suppressed by Tregs. This type of oscillatory responses reflects relapse-and-remission form of autoimmune diseases due to chronic stimulatory factors.

**Figure 2:**
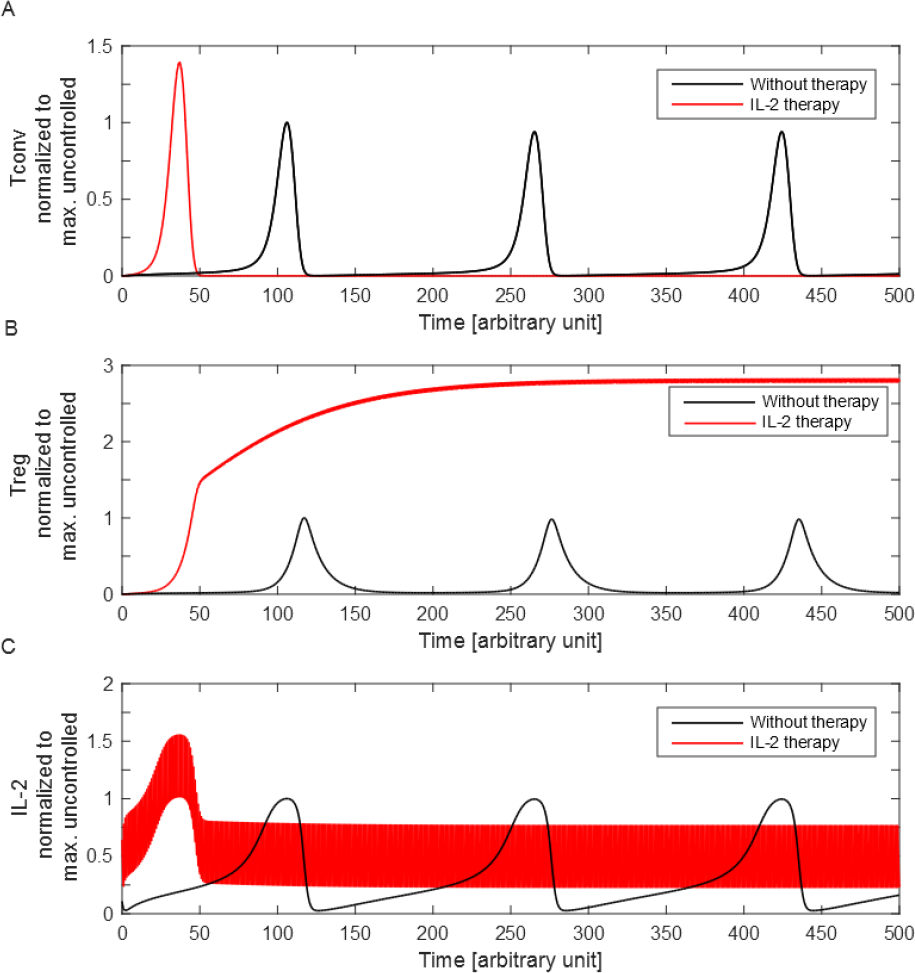
Immune response with and without IL-2 therapy. The T cell response model equation (1) was numerically solved in the presence (red) and absence (black) of IL-2 injections. The values of (A) Tconvs, (B) Tregs and (C) IL-2 concentration were normalized to their maximum value in the absence of IL-2 injections. For IL-2 therapy, constant doses of 0.5 (arbitrary unit) are administered every unit of time (*δ* = 1). The initial conditions are set to zero.

### IL-2 therapy alone cannot avoid acute Tconv response, but controls further relapses

A fixed-dose of IL-2 injections was employed at the time of Ag-stimulation within equidistant intervals. The time evolution of the system variables were obtained in the presence and absence of IL-2 injection (**Figure 2**). With IL-2 therapy, the initial proliferative response of Tconvs starts earlier and reaches a higher peak than without therapy (see **Figure 2A**). The continuation of IL-2 injections keeps the number of Tregs high and prevents re-initiation of Tconv responses.

The undesired consequence of IL-2 therapy in our simulation scenario is the stronger and earlier acute Tconv response. This effect resulted from the augmentation of external IL-2, raising the total systemic IL-2 concentration to a higher value than without the therapy. However, once the number of Tregs increased to a sufficient level, IL-2 injections can maintain a Treg population sufficient to suppress Tconvs. The suppression relies on the continuation of IL-2 injections for as long as the Ag-stimulation persists.

### Restricting acute Tconv responses with adoptive Treg transfer

According to the T cell response model, the IL-2 therapy may control re-initiation of the Tconv response after transiently boosting the first Tconv response. This initial boosting is due to the absence of sufficient Tregs in the early time points needed to suppress Tconvs and compete for IL-2. Therefore, in the context of transplantation where suppression of the acute Tconv response is a necessity for graft accommodation, IL-2 therapy alone may not be a safe immune suppressive strategy. One solution to restrict the rise of the Tconv population in the early episodes of IL-2 injection is to raise the initial number of activated Tregs, by adoptive Treg transfer. This combined strategy is implemented in the numerical simulation by assuming a nonzero initial value for Tregs (i.e. *R*(*t* = 0) = *R*_0_ > 0). Simulations with different level of transferred Tregs were performed to observe the quantitative impact of the therapy on the peak of Tconv responses. A higher amount of adoptively transferred Tregs resulted in a larger reduction of the Tconv peak (**Figure 3A**) and, consequently, a reduced peak of the systemic IL-2 concentration due to less IL-2 secretion (**Figure 3B**). After the IL-2 and Tconv peak, all variables converged to a similar range as determined by the IL-2 dose alone. The simulations differ only in the initial Treg value and, therefore, only a transient impact is induced. The speed of convergence decreases with more adoptively transferred Tregs, which is due to less secreted IL-2 by Tconvs and, thus, lower systemic IL-2 concentrations.

**Figure 3:**
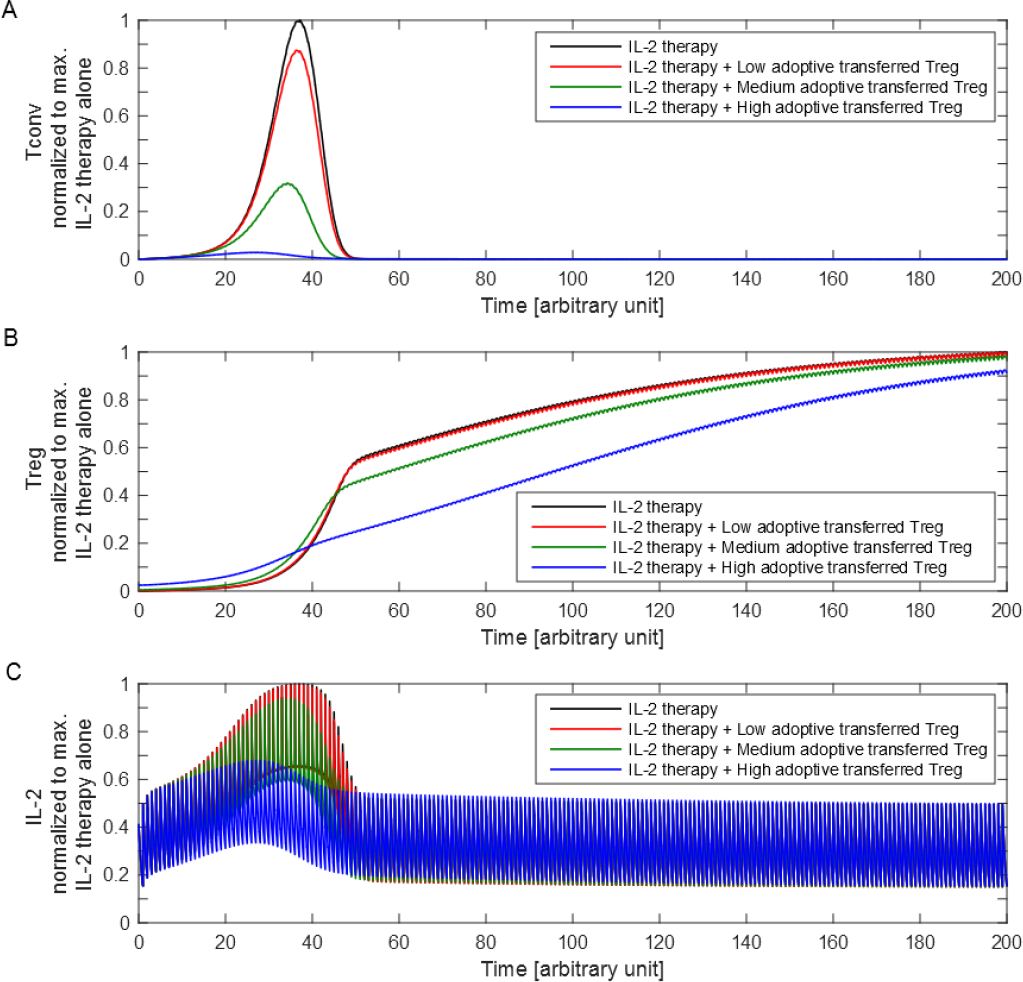
IL-2 therapy combined with adoptive Treg transfer. Equation (1) was solved for the cases of IL-2 therapy alone (black) and in combination with different levels of adoptive Tregs transfer at *t* = 0 (colors). The values of (A) Tconvs, (B) Tregs and, (C) IL-2 concentration were normalized to their maximum value for the case of IL-2 therapy alone. The administration frequency is 1. Low, medium and high adoptive Tregs correspond to initial values of *R* of 0.01, 0.04 and 0.5, respectively.

These results suggest that the combined strategy of IL-2 therapy with an one-time adoptive transfer of Tregs is able to restrict the undesirable effect on Tconv proliferation and the systemic increase of IL-2.

### Fixed versus adaptive IL-2 dosing

So far, IL-2 doses were fixed irrespective of the state of the immune response. In other words, no information of the state of the system was used and, therefore, the knowledge about the interplay between variables of the system is fully neglected. We addressed the possibility of adapting IL-2 doses automatically, taking into account the state of the T cell response at the time of each IL-2 injection as well as pre-defined constraints on the immune variables.

We casted the problem of adaptive IL-2 dosing in the framework of feedback control systems in order to profit from advanced tools in the control engineering discipline. First, at each IL-2 injection, measurements of immune variables (*T*, *R* and *I*) are needed. These measurements are used as an input to the control unit for calculation of the best IL-2 dose with the iZMPC algorithm, as described in the SI. **Figure 1** shows the scheme of the closed loop feedback system between the immune variables and the control unit.

The calculated IL-2 dose is applied to the system at *t* = *τ*_*i*_ (via equation (3)). At the next IL-2 injection (i.e. *t* = *τ*_*i*+1_) the same calculation is repeated. This procedure continues until the variables of the system are steered in the desired ranges (pre-defined constraints). Thus, the IL-2 doses are optimal, in the sense that the control unit proposes the minimum IL-2 dose sufficient to force and keep the variables of the system within the target ranges. The variable IL-2 doses reflect the adaptation of IL-2 doses to the behavior of the system and takes advantage of our knowledge about the immune response as captured in equation (1).

In **Figure 4**, the behavior of the immune variables with fixed versus adaptive dose IL-2 therapy, each combined with adoptive Treg transfer, are compared. While adaptive doses reduced the peak response of Tconvs, the steady state value of Tconvs settled at a higher level and Tregs at a lower level (**Figure 4A-B**). The advantage of the adaptive method becomes evident by noting the kinetics of systemic IL-2 at the time of the Tconv peak response as well as the dose of injected IL-2 (**Figure 4C-D**). The control unit reduced the IL-2 dose when the contribution of Tconvs to IL-2 secretion is increased, as well as at later time points when the Treg population is stabilized. This is also reflected in the lower amplitude of IL-2 oscillations.

**Figure 4:**
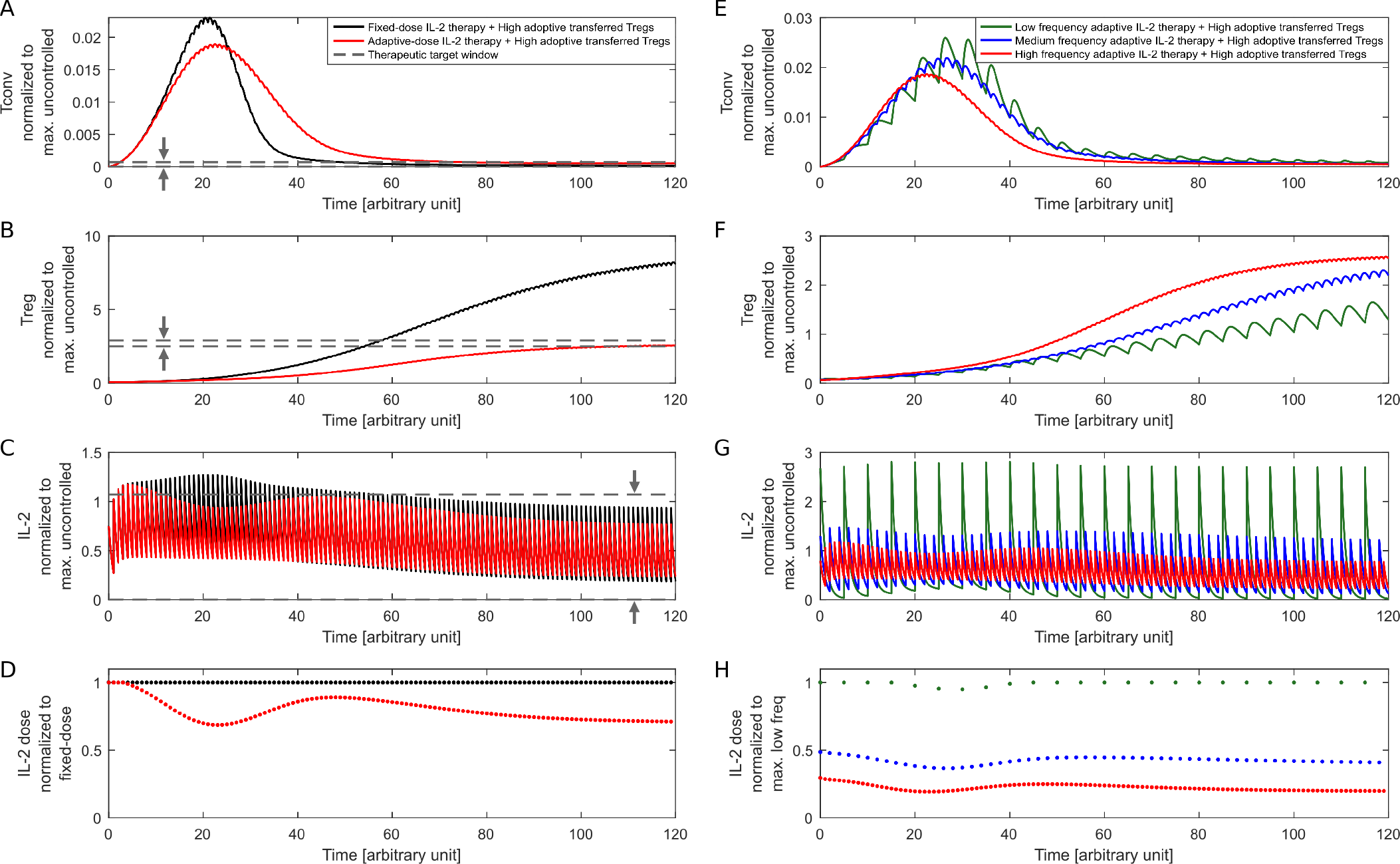
Adaptive IL-2 dosing strategy. The kinetics of (A) Tconvs, (B) Tregs, (C) IL-2, and (D) IL-2 dose is shown for fixed (black) and adaptive dose IL-2 therapy (red), each applied at every time unit (*δ* = 1) and combined with adoptive Treg transfer (*R*(*t* = 0) = 0.5). The physiological range was 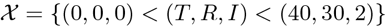 and the therapeutic target window was 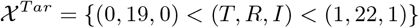. IL-2 doses were constrained to 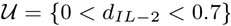. For the fixed dose therapy, the maximum allowed dose (i.e., 0.7) was administered. The calculated adaptive doses successfully enforced variables to the therapeutic target window (horizontal dashed lines). (E-H) Same, assuming different IL-2 injection frequencies *δ* of 1, 2, and 5 time units, corresponding to high, medium and low frequency, respectively. The control unit selects the corresponding suitable doses in each case. The value of Tconv, Tregs and IL-2 is normalized to its maximum value for the uncontrolled case (without IL-2 therapy, Figure 2, black curves). For low frequencies higher doses are required. Therefore, the maximum allowed IL-2 dose was increased to 2.5 to make the optimization problem feasible. The IL-2 dose is normalized to the dose in (D) fixed-dose value, and (H) maximum dose value in low frequency.

These results suggest that the proposed strategy of IL-2 dosing is able to limit systemic IL-2 levels by adaptively reducing the dose when external IL-2 is not needed.

### IL-2 therapy: frequency versus dose

In addition to dose, the frequency of IL-2 injections (*δ*) is a free parameter for designing the therapy. To evaluate the impact of the IL-2 injection frequency on the dynamics of T cells, the frequency was altered and adaptive IL-2 dosing was calculated (see **Figure 4E-H**). In order to control the Tconv response with low frequency IL-2 injections, higher doses of IL-2 are needed. This results in a higher peak of systemic IL-2 concentration.

## Discussion

Feedback control design provides a level of robustness in achieving objectives and constraints in uncertain and complex conditions. It has been successfully applied in engineering applications within multiple aspects of human daily life. Physiology of the human body itself contains many feedback control loops to regulate processes under uncertain and stochastic environmental conditions (32). Feedback control concepts are less common in medicine and therapeutic design. Therapeutic interventions based on feedback allow for adaptation to the unforeseen and unavoidable disturbing factors that are imposed to an individual. Using measurements prior to each therapeutic intervention and evaluating a mathematical model-based prediction of system state-trajectory allows for informed adjustments of the therapy. This approach would lead to a more robust therapeutic outcome than a protocol-based treatments which are typically based on one-fits-all approach. In this study, a novel feedback control scheme inspired from the control engineering field was assessed *in silico* for IL-2 therapy with the aim to regulate T cell responses. The control scheme was applied to a mathematical model of T cell responses that relies on established principles of Tconv and Treg activation, proliferation and regulation (21).

In the context of immune tolerance induction/breakdown, antigen-specific interventions are typically desired. Such interventions require targeting a particular subset of Tconv and Treg clones with high specificity to the antigen. However, IL-2 therapy is an antigen-nonspecific immune intervention that influences all specificities of T cells, and also targets both Tconvs and Tregs. Therefore, violation of tolerable concentration of IL-2 could cause significant dose-related morbidity, such as in application to cancer (33). According to our *in silico* results, a fixed IL-2 dose ignoring the contribution of the endogenously secreted IL-2 would lead to an unwanted increased systemic IL-2 concentration (**Figure 2C**). The adaptive IL-2 dosing scheme could limit this side-effect by taking into account the measured IL-2 concentration prior to injection episodes, as well as enforcing the system to a confined range of IL-2 concentration determined by clinical constraints (**Figure 4C**).

The presented methodology is general and can be adapted to different design requirements, such as increasing Tconv number that would be beneficial in cancer applications. In this study, we targeted tolerance induction with the objective of increasing and stabilizing Treg numbers with specificity to an antigen, such as graft-specific antigen in transplantation. Tconvs and Tregs are both activated by their TCR recognizing the specific antigen. However, once Tregs are activated, their suppressive function is antigen-nonspecific and could suppress Tconv responses against unrelated antigens (34). This bears the risk of unwanted tolerance induction against pathogenic agents (35). Therefore, the converging number of Tregs upon long-term IL-2 treatment and its impact on immunity against other antigens is a therapeutic design concern. In the presented adaptive control scheme, the therapeutic target window of variables can be imposed to the control unit reflecting such clinical constraints (**Figure 1**). As an example, we showed that by confining the Treg numbers to a specific range, even a lower number of Tregs compared to the fixed-dose IL-2 therapy ensures a similar extent of Tconv suppression during the acute phase (compare fixed- and adaptive-dose therapies in **Figure 4A-B)**. Therefore, the proposed control scheme has the flexibility to enforce such clinical constraints into the therapy design (IL-2 dosing).

IL-2 therapy as an antigen-nonspecific approach is often used in conjunction with the antigen-specific ther-apies of adoptive (Tconvs/Tregs) cell transfer with the aim to sustain the vitality and efficacy of the transferred cells (36, 37). The frequency of antigen-specific cells in patients, in particular of Tregs, is typically low and *ex vivo* expansion protocols are required to increase the cell number. In the context of transplantation, our *in silico* results showed that the peak of Tconv response is inversely related to the number of adoptively transferred Tregs that was initially provided to the system (**Figure 3A**), suggesting that increasing the number of adoptively transferred Tregs specific to the graft-antigens increases the chance of graft accommodation. However, prolonged *in vitro* expansion of endogenous antigen-specific Tregs is shown to impair their suppressive function (38). Using chimeric antigen receptors (CARs) to change the specificity of T cells (39) is a promising method to construct a sufficient number of antigen-specific cells. Our adaptive control scheme can incorporate CAR Tregs and then be employed to optimize the IL-2 therapy with the aim to regulate and stabilize their number after transfer to the patient. Note that the adaptive control design is not directly linked to the amount of adoptively transferred cells, as it just temporarily change the initial condition of the system and influences the transient immune response. However, the designed IL-2 dosing changes accordingly to enforce the transient T cell response to the predefined physiological ranges of the system variables (clinical constraints). The long-term state of the system is only dictated by long-term therapies.

Despite attractive benefits of using the presented methodology, its application for an individual patient is challenged by different sources of uncertainty that requires further investigation. The performance of the control design in action relies on the accuracy of the mathematical representation of the system. The mathematical model that we used here contains a low degree of freedom, which simplifies parameter inference from experimental and clinical data. In principle, there is a trade-off between model complexity and parameter identifiability. On the one hand, increasing the model complexity leads to a better representation of the multiple interactions existing in real T cell responses and, thus, increases the control performance. On the other hand, most experimental and clinical measurements are limited by the accessibility of the immune variables, as well as the availability of biomarkers, which may not directly or uniquely be linked to the considered immune variables in the model. This limitation leads to unidentifiability and poor individualization of the model parameters for the patient and, consequently, weakens the control performance. Another source of uncertainty is the environmental or internal disturbing factors that are imposed on the patient during the therapy, such as infections. Due to the nature of feedback control design, the adaptive therapy is calculated according to the current state of the system, and therefore, the therapy would be robust to such disturbing factors. However, such an infection might interfere with the T cell dynamics of the system, which is not reflected in the current model. The employed mathematical model represents a monoclonal T cells response. The impact of the IL-2 therapy on polyclonal or concurrent T cell responses needs further investigations.

We provided qualitative *in silico* results for an adaptive control scheme in IL-2 therapy, as a proof-of-principle to motivate further investigations in this direction. There are more and more promising results from IL-2 therapies in experimental studies and clinical trials, which calls for an interdisciplinary approach to bring the presented methodology to a quantitative level and pave the way for ultimate incorporation and validation in translational studies and clinical trials.

## Authors contributions

SK designed the study. GM and SK proposed the adaptive control scheme. GM developed the adaptive control methodology and performed the simulations. MHH supervised the study. All authors contributed to the interpretation of the results and writing the manuscripts.

## Acknowledgements

SK was supported by the German Federal Ministry of Education and Research within the Measures for the Establishment of Systems Medicine, project SYSIMIT (BMBF eMed project SYSIMIT, FKZ: 01ZX1308B) and by the Helmholtz Association, Zukunftsthema Immunology and Inflammation (ZT-0027).

## Supplemental Information (SI)

### Algorithm of adaptive IL-2 dose calculation

In the following, the steps toward calculation of adaptive doses of IL-2 using iZMPC are explained. Consider the nonlinear system

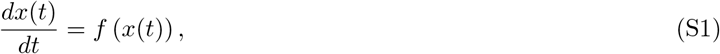

where *x* ∈ ℝ^*n*^ is the vector of system dynamics (here, *x* = [*T, R, I*]). Suppose *u* ∈ ℝ is the drug dose (here, *d*_IL−2_) which affects the system at the discrete time intervals *τ*_*i*_, i = 1, 2, by sudden changes in the state variables

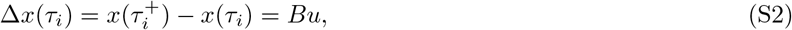

where 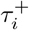 denotes the time instant after *τ*_i_. *B* ∈ ℝ^*n*^ models the impact of *u* on the states, and the amplitude of the sudden pulses at *τ*_*i*_ is equal to *Bu*. We assume equidistant pulses, i.e., *τ*_i+1_ − *τ*_*i*_ = *δ*, *i* = 1, 2,…. Thus the full system is modeled in the template of nonlinear impulsive systems (22) as an augmentation of equations (S1) and (S2).

Depending on the considered biological framework, different constraints may arise; e.g., drug doses are constrained within the physiologically approved limits and also states should be kept within their functional regions. With

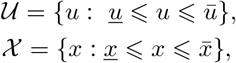

and an arbitrary target set 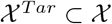 (therapeutic target window), the aim is to compute 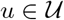 to force *x* moving from its initial value *x*(0) to a point in 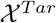. Calculation of *u* is based on the iZMPC (23). In what follows, we delineate the preliminary steps toward using iZMPC. A detailed description, the mathematical basis of the steps and some other biological application of iZMPC can be found in (19, 20, 24, 25).

#### Step 1: finding equilibrium points (*x*_*s*_, *u*_*eq*_)

Augmented system of (S1) and (S2) can be reformulated as 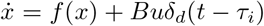 where *δ*_*d*_(*t* − *τ*_*i*_) is the Dirac delta function

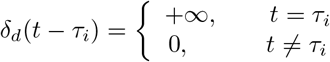

Assume continuous delivery of the drug and calculate 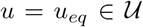 and *x*_*s*_ satisfying the steady state condition *f* (*x*_*s*_) + *Bu*_*eq*_ = 0.

#### Step 2: finding equilibrium levels (*x*_*s*_, *u*_*s*_)

Find *u* = *u*_*s*_ such that the impulsive system (augmented equations (S1) and (S2)) with Δ*x* = *Bu*_*s*_ and impulse frequency *δ* reaches almost the same equilibrium level as *x*_*s*_. Note that, different *δ* result in different *u*_*s*_.

#### Step 3: linearization

Calculate 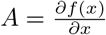 at *x* = *x*_*s*_.

#### Step 4: shift constraints

Calculate shifted sets 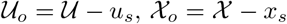 and 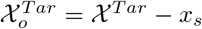.

#### Step 5: feasible generalized control equilibrium zone (set)

Compute two new sets 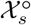 and 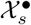 such that

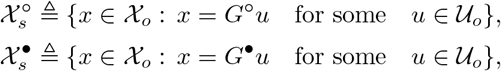

where

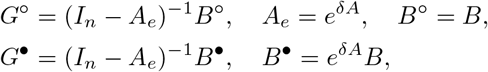

and *I*_*n*_ is the identity matrix of dimension 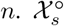 and 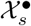 implicitly generate the input equilibrium set

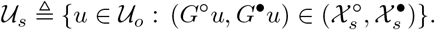

#### Step 6: generalized equilibrium zone (set)

Compute 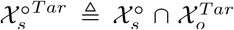 and 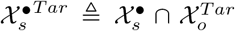. Correspondingly, we can obtain 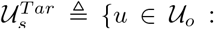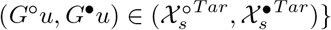. If 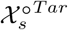 or 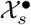 is empty, the control problem is not properly formulated and 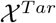 must be increased or *δ* should be decreased. There is a free set computation toolbox “mpt3” in MATLAB which can be downloaded at http://people.ee.ethz.ch/mpt/3/.

#### Step 7: MPC input

At each *t* = *τ*_*i*_, we use the current state of the system of augmented equations (S1) and (S2) *x* and provide *x* − *x*_*s*_ as input to the iZMPC algorithm (see Step 8) which determines *u*.

#### Step 8: iZMPC problem

MPC is a finite time-horizon optimization problem which receives the current state of the system and returns U = {**u**(0), **u**(1), …, **u**(*N* − 1)} (with *N* the control horizon). It predicts the next *N* states of the system using the sampled current state and calculates the next *N* control actions (here, IL-2 doses). Only the first calculated input, i.e. **u**(0) is applied to the system and this process is repeated at every sampling time.

iZMPC is an MPC which at each impulse *τ*_*i*_, *i* = 1, 2, … (i.e., the sampling times) takes *x*(*τ*_*i*_) and calculates U for the impulsive system. Note that, iZMPC is mainly developed for linear impulsive systems. Using the method of linearization around equilibrium levels makes it possible to apply iZMPC to linearized impulsive systems which are originally nonlinear. In the case that errors due to linearization are not acceptable, one may have to stretch out for nonlinear impulsive MPC (20).

The optimization problem to be solved at each *τ*_*i*_ by iZMPC is given by

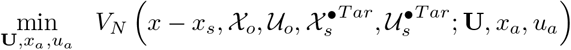

subject to

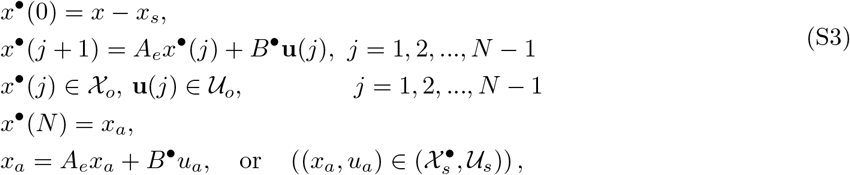

where

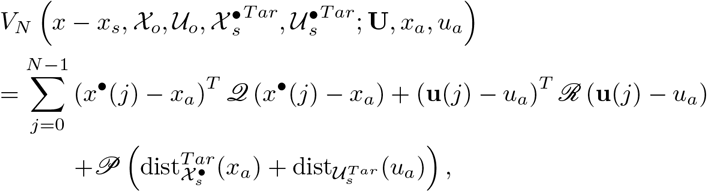

and 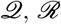 and 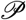 are positive definite matrices and positive numbers respectively. The transit behavior of the system under iZMPC can be tuned using these weighting matrices and parameters. In addition, 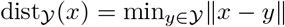.

Note that, 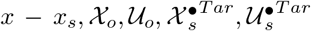 are given parameters in the optimization problem, whereas U = {**u**(0), **u**(1), …, **u**(*N* − 1)}, *x*_*a*_ and *u*_*a*_ are the optimization variables. When the iZMPC problem (S3) is solved, the optimal drug dose *u* in the system of augmented equations (S1) and (S2) (or *d*_IL−2_ in (3)) is obtained by *u* = **u**(0) + *u*_*s*_.

